# Sex-biased duplicates are rapidly generated during *Drosophila* tRNA repertoire evolution

**DOI:** 10.1101/2025.11.24.689867

**Authors:** Dylan Sosa, Marek Sobczyk, Jianhai Chen, Shengqian Xia, Tao Pan, Manyuan Long

## Abstract

Eukaryotic genomes encode hundreds of transfer RNA genes ostensibly to ensure efficient translation. How these seemingly redundant tRNA copies arise and are naturally selected in metazoan genomes and populations remains largely unexplored, owing to challenges in accurate sequencing of tRNA loci and their transcripts. We leveraged long-read genome assemblies of 24 Drosophilid species to infer the origination times of the entire *Drosophila melanogaster* tRNA gene repertoire and found that continuous gain and loss of tRNA duplicates throughout 60 million years of divergence has resulted in rapid taxonomic restriction of isodecoding and isoaccepting copies––even producing one isodecoder specific to *D. melanogaster*. Moreover, we identified patterns of global codon usage, especially in lineage-specific genes, incongruous with translational efficiency hypotheses. Through generation of tRNA sequencing in *D. melanogaster* we observed that recently duplicated, taxonomically restricted tRNA copies had sexually dimorphic patterns of expression, fragmentation, and nucleoside modification. Our work implicates the emergence of taxonomically restricted tRNA genes as sources of regulatory diversity and reveals that sexual and natural selection affect the evolutionary dynamics of the tRNA gene family in metazoan species.

## Main Text

Chiefly known for their central role in protein synthesis through mRNA translation, transfer RNAs constitute one of the largest non-coding gene families in all domains of life. Most tRNA genes are encoded by the nuclear genome and are organized into isoaccepting groups (isoacceptors) based on their anticodon sequence which specifies the amino acid they are charged with by cognate aminoacyl-tRNA synthetases. Isoaccepting groups can be further subdivided into isodecoding groups (isodecoders) based on substitutions outside of the anticodon loop, furthering the diversity of the tRNA gene family (*1*). Eukaryotic species particularly harbor large and seemingly redundant tRNA gene repertoires––the *Homo sapiens* genome encodes 416 tRNA genes while *Saccharomyces cerevisiae* encodes 275 (*2*). However, little is understood regarding how identical and nearly identical nuclear tRNA gene copies proliferate and are maintained in eukaryotic genomes and populations.

While the expansion of nuclear encoded tRNA gene pools has been of long-standing general interest, studies have often focused on codon optimality (*3, 4*) or the primordial origins of tRNA structure (*5, 6*). Studies of the evolutionary history and principles that shape tRNA gene complements in eukaryotes have been constrained by the highly similar and short length of tRNA genes, a lack of syntenic comparison of tRNA loci between multiple long-read assemblies, and difficulties in high-throughput quantification of tRNAs caused by post-transcriptional base modifications (*7–10*).

The substantial variation in isoacceptor type and number between closely related organisms is generally thought to reflect a co-evolutionary relationship between tRNA availability and mRNA demands (*11, 12*), but this balance may be less straightforward in specific tissues or disease states in metazoans (*13*). Given the significant interspecies variability of tRNA gene copy numbers and their critical role in mRNA decoding, we investigated *Drosophila* tRNA gene origination through analyses of long-read genome assemblies and generation of small RNA sequencing. We investigated how selective forces maintain new duplicates, how the diversity of eukaryotic tRNA repertoires is influenced by new coding genes, and what role sex has in the differential emergence or function of tRNA duplicates.

## RESULTS

### Taxonomically restricted genes exhibit unbiased codon patterns incongruous with nuclear-encoded tRNAs

Adaptive expansion of the cytosolic tRNA pool through duplication may be an effective strategy to ensure mRNA decoding rather than codon optimization of every gene through point mutations (*14, 15*). Despite limited phylogenetic conservation, new genes acquired through duplication and other mechanisms (*e.g.* retrotransposition, *de novo*) often have biased expression in reproductive and neural tissues (*16, 17*), integrate into conserved pathways (*18*), and significantly affect organismal fitness (*19, 20*). As evolutionarily young genes arise in metazoans some may be unoptimized in their codon usage (*21*), potentially resulting in discordance between their mRNA and the available pool of tRNAs. To test if tRNA repertoires are generally coevolving to adaptively reflect global codon usage, we first measured the relationship between biased synonymous codon usage, ψ*_g_*, and the tRNA adaptation index, *tAI_g_*, of 13,785 *D. melanogaster* protein coding genes divided into two groups based on their age (young < 40Ma, ancient ≥ 40Ma) (*22*). We observed significant correlations (*p* < 0.02 and *p* < 0.001, Pearson’s correlation) between codon usage and adaptation to the genomic tRNA pool for both young (*n* = 843) and ancient genes (*n* = 12,942), with a reduction in both the correlation coefficient *S* and the coefficient of determination *R*^2^of young genes compared to ancient (Fig. 1A,B). We next examined genes stratified by their age and inferred origination mechanism: DNA-based duplication, RNA-based duplication, or lineage-specific orphans (*23*). Ancient genes across mechanism types had significant, positive correlations between ψ*_g_* and *tAI_g_* (*p* ≤ 0.004, Pearson’s correlation) as well as elevated *R*^2^ compared to young genes of each mechanism type. There was no observed trend between ψ*_g_* and *tAI_g_* amongst young lineage-specific and RNA-based duplicate genes (*p* = 0.701 and *p* = 0.0617, Pearson’s correlation), but the codon bias of DNA-based duplicates was significantly correlated (*p* = 0.0331, *R*^2^ = 0.01). In both age groups lineage-specific genes had the weakest *S* and *R*^2^ values (*S* ≤ 0.28, *R*^2^ ≤ 0.08), emphasizing that despite the proliferation of tRNA genes within eukaryotic genomes translational selection is not strongly affecting codon composition in recently derived genes or those putatively originated from intergenic regions (Fig. 1B, C).

**Fig. 1.**
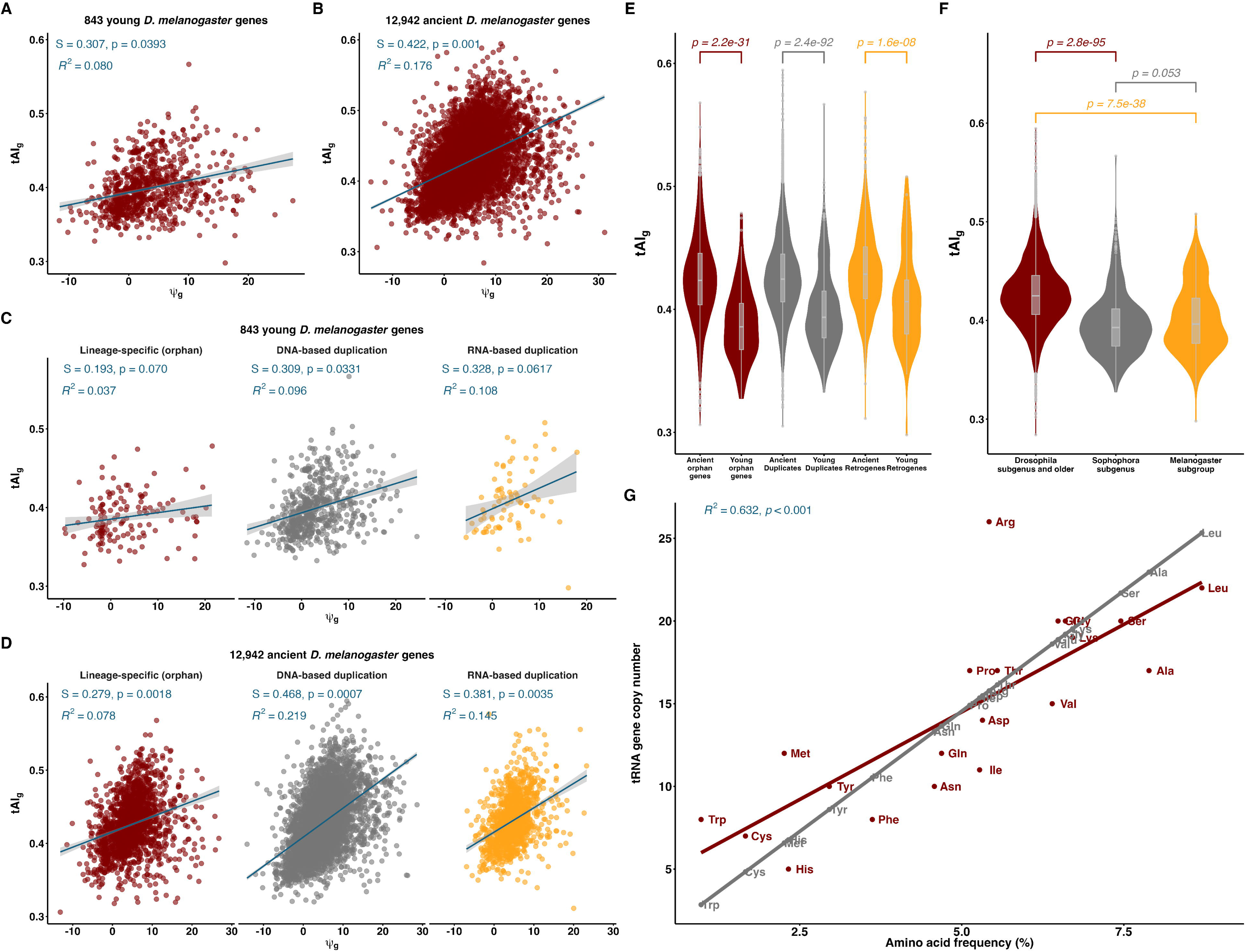
Recently derived *D. melanogaster* genes are poorly adapted to the nuclear encoded tRNA pool. **(A** and **B)** Test of translation selection using the Pearson correlation (*S*) of biased synonymous codon usage values, ψ_*g*_,and the tRNA adaptation index, *tAI*_*g*_, of (A) 843 young and (B) 12,942 ancient *D. melanogaster* protein coding genes. Empirical p-values determined by simulation and permutation. **(C** and **D)** Same as (A, B) but show lineage-restricted (young) genes and conserved (ancient) genes divided by their inferred origination mechanism. **(E** and **F)** Wilcoxon rank-sum test of the distribution of tRNA adaptation index, *tAI*_*g*_, for 13,785 *D. melanogaster* protein coding genes. (E) Distribution of *tAI*_*g*_ by mechanism of gene origination. (F) The same as (E), but showing genes divided into three age groups in order of earlier divergence to most recent: *Drosophila* subgenus-specific and older, *Sophohora* subgenus-specific, and *melanogaster* subgroup-specific. **(G)** Correlations of observed tRNA copy numbers (red) and expected tRNA copy numbers (gray) with usage frequencies of their corresponding amino acids (*24*).

We then compared the distributions of young and ancient gene *tAI*_g_ values for each origination mechanism to interrogate whether unbiased synonymous codon usage was a general pattern of new gene origination. For each comparison (Fig. 1D) we observed a significant reduction in *tAI*_*g*_ values of young genes compared to the distribution of ancient genes (*p* ≤ 1.6 × 10^-8^, rank-sum test). This trend was again observed when splitting the *D. melanogaster* genes into three age groups in order of most distant to most recent divergence time: *Drosophila*-specific and older, *Sophophora* subgenus-specific, and *melanogaster*-subgroup specific. The two younger age groups each had significantly reduced *tAI*_*g*_values compared to the *Drosophila*-specific distribution (*p* = 2.8 × 10^-95^ and *p* = 7.5 × 10^-38^, rank-sum test) and there was a non-significant difference between *Sophophora* subgenus and *melanogaster*-subgroup values (*p* = 0.053, rank-sum test) (Fig.1E), illustrating a downward tendency of *tAI*_g_as gene age decreases. Although a significant correlation (*R*^2^= 0.632, *p* < 0.001) between amino acid usage frequency and *D. melanogaster* tRNA copy numbers was observed (*24*), all isotype copy numbers were distinct from values expected by amino acid usage (Fig. 1G).

Together, these results reveal that codon usage of recently evolved genes regardless of origination mechanism are maladapted to the nuclear encoded tRNA repertoire, and that tRNA copy numbers exceed or do not meet those predicted by translational efficiency hypotheses. This suggests nuclear encoded tRNA copy numbers are not generally adaptively coevolving with global codon usage (Fig. S2A-C). Rare codons, those non-optimal for translation efficiency, can be necessary for functions of recently derived proteins (*25*) and post-transcriptional remodeling of tRNAs could be a preferred strategy to meet codon usage patterns rather than adaptive duplication (*26, 27*). Thus, the tRNA gene family in *D. melanogaster* may be expanding under pressures besides selection for efficient decoding of mRNA.

### tRNA gene repertoires rapidly evolve in copy number and isotype

Although duplication is known to be a major mechanism by which tRNA genes arise, little is known regarding tRNA gene turnover in closely related species, their duplication patterns, or how tRNA repertoires are maintained in metazoan genomes and populations (*28*). We first annotated nuclear-encoded tRNA gene sets of 24 Drosophilid species long-read assemblies (Table S1) with tRNAscan-SE v.2.0.9 (*29*) and observed highly variable tRNA copy numbers between the 24 closely related species considered in this work (Fig. S1A - C). Total Drosophilid tRNA gene content ranged between 241 to 428 genes and unique isodecoding sequences ranged between 88 to 165 per species (Fig. 2A). In our focal species, *D. melanogaster*, we annotated 291 nuclear-encoded tRNA genes of which 89 are unique isodecoding sequences. Following these copy number observations, we hypothesized that tRNA gene turnover may occur more frequently in metazoans than expected despite their importance to the central dogma. To test this, we developed a gene age dating methodology to infer the temporal distribution of *D. melanogaster* tRNA gene origination.

**Fig. 2.**
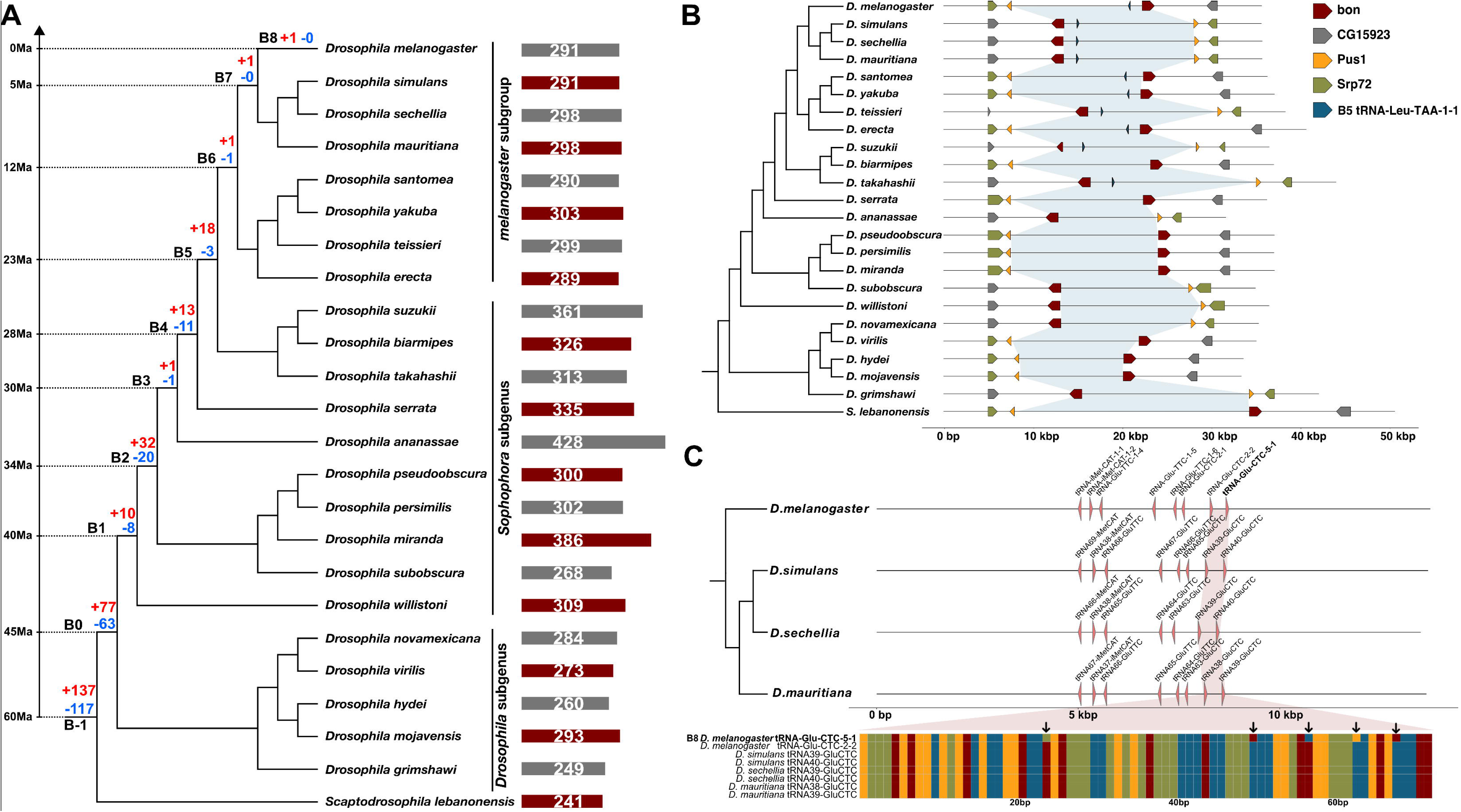
tRNA gene origination is a dynamic process. **(A)** Drosophilid species divergence time phylogeny. See references for divergence estimation (*32, 33*). Values in red indicate the number of *D. melanogaster* tRNA gene births at a given branch while blue values represent the total number of loss events of genes originating from the previous branch. **(B)** Schematic illustrating the gene age dating method applied to Drosophilid genomic tRNAs. For example, we inferred the time of origination of the young isodecoder *tRNA-Leu-TAA-1-1* as occurring at branch 5 (B5), approximately 23Ma ago, as the earliest diverging species containing a copy of this gene is *D. takahashii* in the clade just before the divergence of the *melanogsaster* subgroup. Blue shaded area represents the conserved syntenic area used for homology detection. **(C)** Multiple sequence alignment of eight tRNA^Glu^ genes from four species in the *melanogaster* subgroup highlighting the sequence divergence of the *D. melanogaster*-specific isodecoder *tRNA-Glu-CTC-5-1*.

As tRNA genes are short, largely identical, and often tandemly duplicated, standard phylogenetic or whole genome alignment strategies to determine gene age may be confounded by ambiguous alignments and imprecise homology detection when attempting to investigate their evolution (*30*). Our age dating strategy reduced the homology search space and probability of incorrectly aligning tRNA genes by determining syntenic regions between conserved, neighboring “anchor” protein coding genes. We assigned each *D. melanogaster* tRNA gene to a branch on the Drosophilid divergence time phylogeny (*31–33*) numbered between -1 and 8 (B-1 through B8, larger values indicate more recent divergence) corresponding to the inferred time of their duplication (Fig. 2B). We observed the origination of *D. melanogaster* tRNA genes to be a continuous process occurring at a rate of 1.93 gene births per Ma (77 genes / 40 Ma). We found that 26.46% of the tRNA gene repertoire is restricted to the *Sophophora* subgenus (B1 – B8) and 1.03% is further restricted to the *melanogaster* subgroup (B6 - B8). We identified one *D. melanogaster*-specific tRNA isodecoder, *tRNA-Glu-CTC-5-1*, which contains five fixed substitutions compared to outgroup species (Fig. 2C), highlighting rapid substitution as previously reported at tRNA loci (*34*).

The fixation of gene loss is also an important contributor to genomic and phenotypic evolution (*35*). For a given *D. melanogaster* tRNA gene we determined the presence or absence of an orthologous, syntenic tRNA in species which diverged after the inferred time of origination of the focal gene. We estimated the rate of tRNA gene loss for genes originating in *Sophophora* subgenus and *melanogaster* subgroup species (B1 – B8) to be 1.1 tRNA genes per Ma (44 genes / 40 Ma). In *D. melanogaster* we observed that 21% (62/291) of the tRNA gene repertoire is not lost in any species following their emergence, possibly constituting a “core” set of tRNA genes as has been proposed previously (*36*). The anticodons of those tRNAs correspond to 16 genetically encoded amino acids (Data S2). This suggests some tRNA duplicates primarily perform their canonical role in translation while others that are lost or gained may be free to acquire “forbidden” substitutions as postulated by Ohno facilitating the acquisition of copy-specific, non-canonical functions (*37, 38*). Of those tRNAs that are retained in all species considered in this work, 32 are unique isodecoding sequences which represent 36% of all *D. melanogaster* isodecoder subfamilies. This suggests increased nucleotide diversity through isodecoder substitutions may allow non-exact duplicates to be maintained over long periods of evolutionary time beyond speciation events. Thus, while mRNA translation is a deeply conserved process tRNA gene repertoires are dynamic and rapidly evolve in copy number and isotype over short evolutionary timescales.

### Lineage-specific tRNA duplicates demonstrate sex-biased expression and fragmentation

As sexual and sexually antagonistic selection have been found to be driving forces in the emergence of new protein coding genes, we hypothesized that sex-related selection could underlie tRNA duplication or differences in their abundance or modification (*39*). To comprehensively interrogate whether sex or gene age affect the expression of tRNAs, we dissected gonad tissues (ovary, testis) from Iso-1 *D. melanogaster* adults in biological triplicate and performed multiplexed small RNA sequencing of total RNA extractions (MSR-seq, Data S3) (*40*). We observed sexually dimorphic abundances of isodecoders and individual isoaccepting duplicates that originated at different times along the *Drosophila* phylogeny (Fig. 3A,B). One of the youngest tRNA isodecoders we identified previously, *tRNA-Lys-TTT-2-5* which duplicated in the *melanogaster* subgroup at B7 just prior to the divergence of *D. melanogaster*, was significantly upregulated in the ovary (Fig. 3C) as a precursor tRNA transcript (pre-tRNA, *p* < 0.05) suggesting sex-specific, non-translational functions (*41*). Moreover, the next youngest differentially expressed tRNA, *tRNA-Asp-GTC-3-1* which we inferred to have originated in the *Sophophora* subgenus at B5 prior to the divergence of the *melanogaster* subgroup, is upregulated in the testis as an antisense transcript (*p* < 0.001) and may perform male-specific signaling or transcriptional regulatory functions through novel interactions with mRNAs (*42*). In total, we identified 38 tRNA genes spanning B-1, B0, B1, B2, B5, and B7 that are differentially expressed between the sexes. These upregulated tRNA transcripts represent 10 isoaccepting families and are primarily antisense or pre-tRNA, suggesting sexual selection not of mRNA decoding efficiency, but of extra-translational functions of tRNAs or their fragments sufficient to maintain redundant isoacceptors and isodecoders in the shared genome (Data S3).

**Fig. 3.**
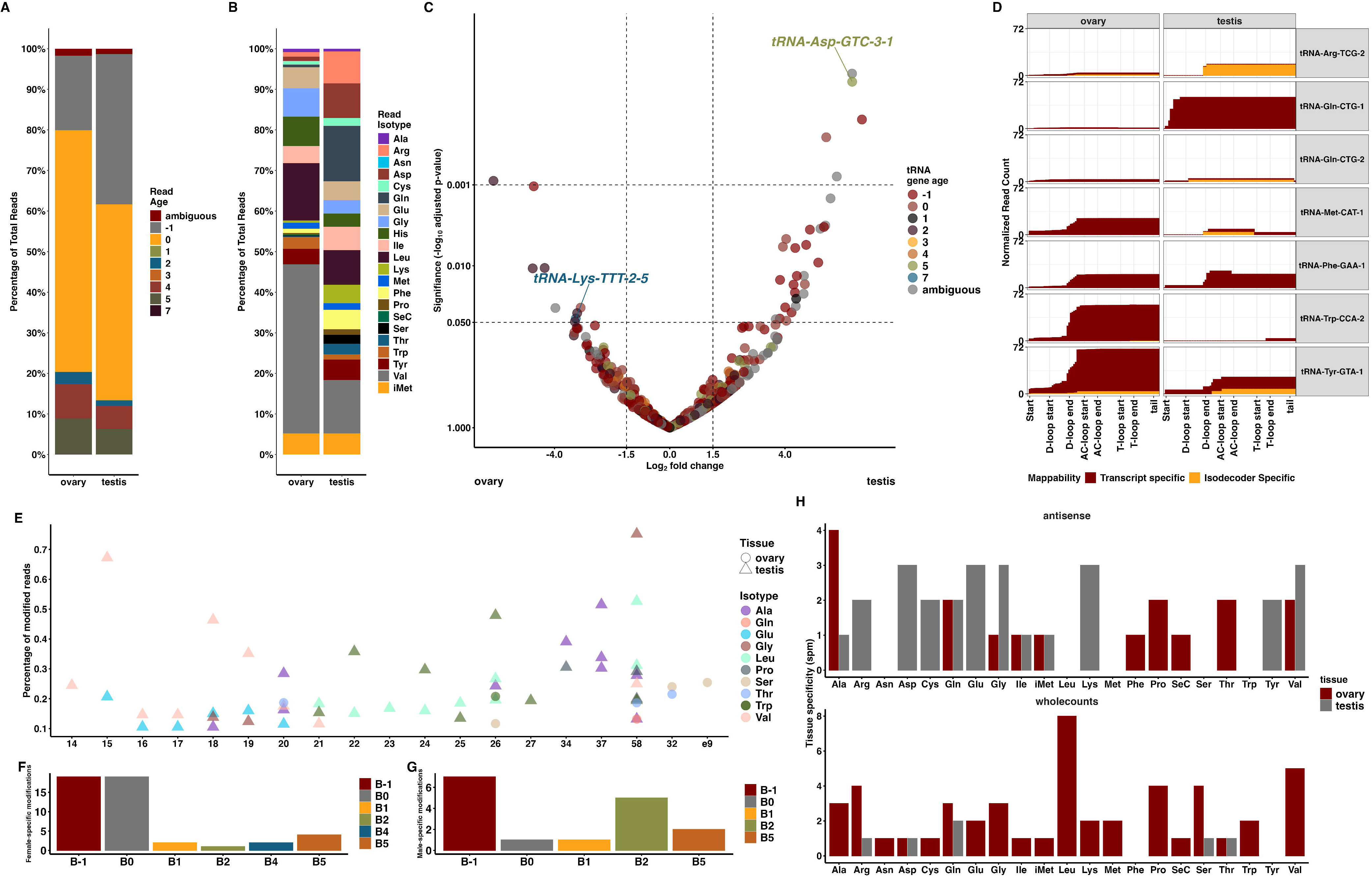
The *D. melanogaster* tRNA repertoire exhibits sex-specific expression, fragmentation, and modification. **(A** and **B)** tRNA sequencing read distributions between *D. melanogaster* gonads. (A) Proportion of tRNA-seq reads colored by inferred gene age and (B) distribution of reads colored by tRNA read isotype. **(C)** Volcano plot of differentially expressed tRNAs genes between ovary and testis (rlog normalization, using DESeq2). Points are colored by gene age unless there is ambiguity due to identical tRNA sequence. **(D)** Bar plots showing the start and end positions of ovary- and testis-specific patterns of tRNA fragment abundance colored by specificity of reads. Red bars indicate reads specific to an isotype; yellow bars are isodecoder specific reads. **(E)** Scatter plot of post-transcriptional modifications specific to either sex. Circles indicate ovary-specific modified transcripts while triangles indicated testis-specific modified transcripts. **(F** and **G)** Bar plots illustrating counts of sex-specific post-transcriptional modifications colored by gene age. (F) Ovary-specific modifications. (G) Testis-specific modifications. **(H)** Bar plots of tRNA-seq reads with biased tissue specificity (spm ≥ 0.9) where red indicates ovary reads and gray indicates testis (antisense above and whole tRNA below).

In addition to abundance, we asked whether tRNA fragmentation, an increasingly appreciated source of tRNA functional diversity is also variable between sexes (*43*). We observed tissue- and isodecoder-specific differences in fragmentation patterns by determining the start and end position of fragmented transcripts (Fig. 3D). Overall, we observed 43 isodecoders as the source of sexually dimorphic fragments (Data S4) between *D. melanogaster* reproductive tissues. Identical tRNAs exhibited sex-specific fragmentation patterns such as *tRNA-Trp-CCA-2,* which is highly upregulated in ovary, but lowly expressed as a 3’ fragment in testis; *tRNA-Gln-CTG-1* which is upregulated in testis but not ovary, *tRNA-Tyr-GTA-1* which is expressed in both tissues, but is biased toward the ovary; and *tRNA-Phe-GAA-1* isoacceptors which are equivalently expressed in both sexes, save for a fragment consisting of the D-loop end through the anticodon loop end. Further, we identified instances of isodecoder and sex-specific fragmentation such as *tRNA-Met-CAT-1, tRNA-Arg-TCG-2,* or *tRNA-Gln-CTG-2* which have testis-limited expression exclusively from those isodecoding sequences and not others of the same isotype, underscoring the functional divergence of nearly identical isodecoding tRNAs between reproductive tissues and highlighting the importance of sex in understanding tRNA duplication.

As post-transcriptional modification of tRNAs is essential for their maturation, function, regulation, and conformational rigidity (*44, 45*), we asked if sex or gene age are associated with certain modifications. We tested this by characterizing the distribution of base modifications along the lengths of tRNA transcripts, and we focused our analyses on modified nucleosides found exclusively in either sex which we observed at positions: 14–27, 32, 34, 37, 58, and e9 (Fig. 3E). Age distributions of tRNAs with sex-specific modifications were similar between the ovary and testis, but ovary-specific base modifications occurred primarily on ancient tRNAs (Fig. 3F, G). Most sex-specific modifications identified were female-biased, suggesting unique tRNA remodeling is necessary to meet translational or extra-translational requirements of the ovary. Strikingly, we observed that young isodecoding genes such as *tRNA-Trp-CCA-2* and *tRNA-Trp-CCA-1*, which originated at B5 after the divergence of the *melanogaster* subgroup, exhibit sexually dimorphic post-transcriptional modification at G26, likely m^2^G26 or m^2^ G26, respectively (*46*).

To investigate whether tRNA tissue-specific expression is generally correlated with gene age, as has been reported for recently evolved protein coding genes (*47, 48*), we computed the tissue specificity measure (spm) (*49*) for all tRNA transcripts. We focused our analysis on those trancripts which had spm ≥ 0.9, indicating strong specificity, resulting in 412 tissue-biased transcripts (Fig. 3H). We divided them into three groups as previously (*Drosophila*-specific and older, *Sophophora* subgenus-specific, and *melanogaster*-subgroup specific) and observed that ovarian tissue consistently had more strongly tissue-biased tRNA transcripts than testis regardless of gene age (*p* ≤ 0.0034, rank-sum test, Fig. S3). However, when discriminating by type we found that antisense transcripts across isoacceptor groups were biased toward the testis, again suggesting possible inhibitory functions necessary to modulate the transcriptomic complexity of the testis which is uncorrelated with other tissues across metazoans (*50, 51*). The opposite was observed for whole tRNA transcripts which were observed to be primarily specific to the ovaries. These data suggest gene age is related to the rate of tRNA base modification and their functional potential, possibly promoting sex-limited neofunctionalization of new duplicates or their fragments. Thus, selection of sex-biased expression may also underlie tRNA repertoire evolution and their functional diversification.

### Recently duplicated isodecoders are selectively maintained in natural populations

Following our observations of tRNA genes originating at various timepoints during *Drosophila* evolution we hypothesized that Darwinian selection would be accelerating the differential emergence and retention of tRNA genes amongst eukaryotic species. We leveraged population genetic data of *D. melanogaster* to test for interpopulation selection signals using the site frequency spectra (SFS) of genomic windows containing tRNA loci. We retrieved 727 accessions of *D. melanogaster* natural population samples representing the putative ancestral range in Africa (AFR) and locations outside of Africa (OOA) (*52*). We computed nucleotide diversity, *π*, and the fixation index, F*_st_*, in 400bp sliding windows with a 50bp step size along the *D. melanogaster* genome. We reasoned that if tRNA genes are continuously duplicating such that interspecific tRNA copy number and isotypes diverge drastically, then they would exhibit significant nucleotide diversity between *D. melanogaster* populations indicative of natural selection. By comparing the ratio of *π*_*+,_ and *π*_--*_ to the weighted F_()_ of tRNA loci we identified six candidates: B-1 genes *tRNA-Lys-CTT-1-5*, *tRNA-Gly-TCC-1-1*, *tRNA-Ser-CGA-1-1,* and *tRNA-Trp-CCA-2-6;* B2 *tRNA-Arg-TCG-2-1,* and B4 *tRNA-Lys-CTT-1-11* (Fig. 4A). Genomic windows containing each of these six tRNA genes had significant reductions of *π* in African populations and increased F*_st_*, suggesting some isodecoding copies may be adaptively maintained in distinct populations of *D. melanogaster*.

**Fig. 4.**
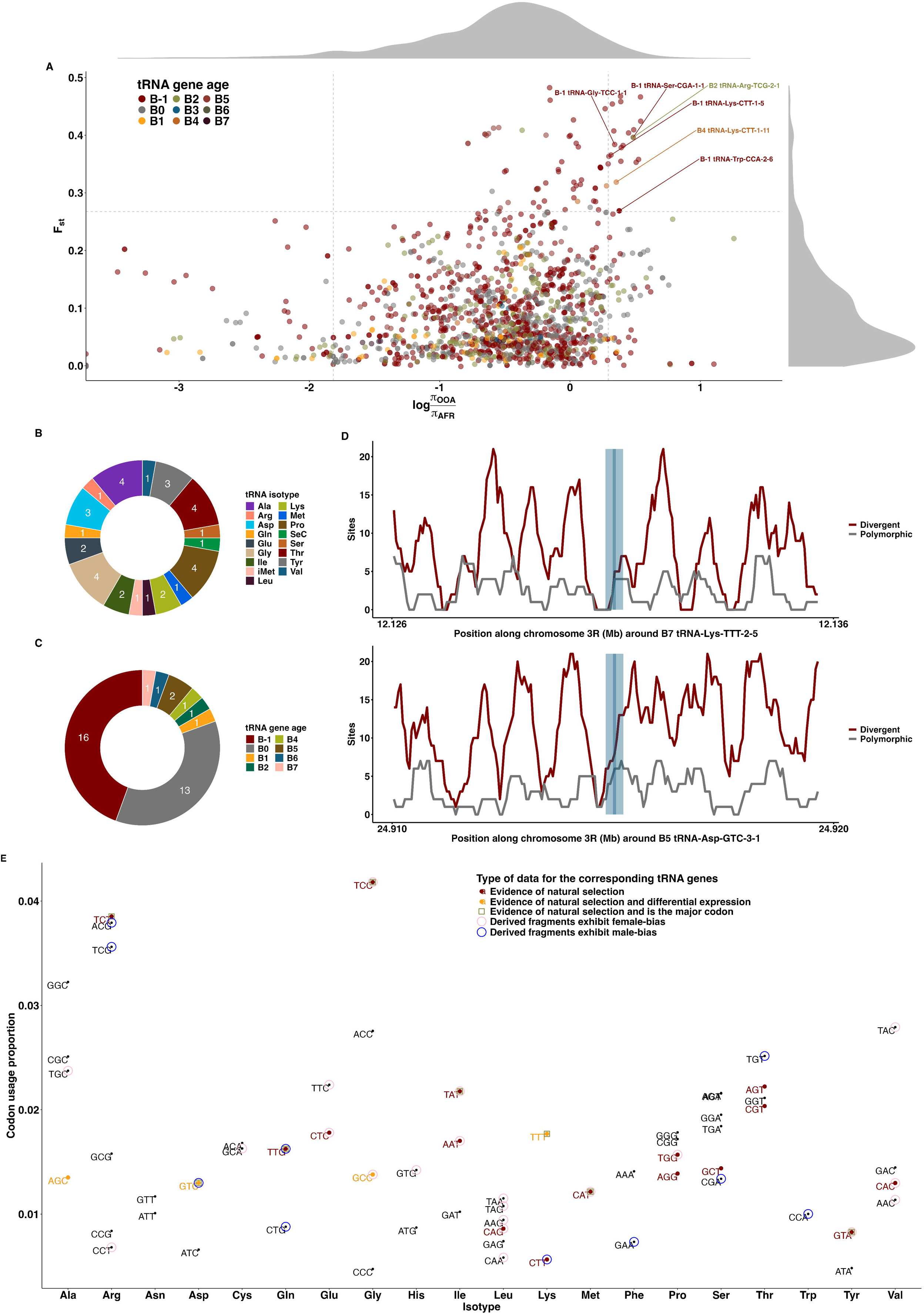
Natural selection maintains tRNA genes but does not necessarily cause repertoire expansion. **(A)** Log ratio of tRNA nucleotide diversity (*π*) between African (AFR) and outside of Africa (OOA) population samples plotted against the weighted fixation index (F*_st_*). Gray dashed lines indicate the significance thresholds used (F_()_ = 2.68, 95th = 2.93, 5th = −1.82). The six windows containing significant tRNA genes are labeled in the upper right quadrant. **(B** and **C)** 36 tRNA genes are within genomic regions with significant HKA-like tests results. (B) 17 isotypes are represented by the naturally selected tRNA genes. (C) Gene age distribution for the 36 significant tRNA genes. **(D)** Two recently duplicated tRNAs (B7 and B5) *tRNA-Lys-TTT-2-5* and *tRNA-Asp-GTC-3-1* are in regions with significant reductions in proportions of segregating to divergent sites polarized with *D. simulans* and *D. yakuba* as compared to neutral expectations (dark blue is the gene region, light blue is the 400bp window centered on the gene). **(E)** Anticodons of each tRNA isotype are plotted according to their frequency of usage in 12, 942 *D. melanogaster* protein coding genes. Anticodons colored in red indicate significant Hudson-Kreitman-Aguadé-like test values (Data S6), while those colored in yellow are differentially expressed in addition to having significant HKA-like results. Those anticodons marked by a green box have signatures of natural selection and are the major codon of a given isotype determined by codon usage frequency. Blue and pink circles indicate sex-biased patterns of tRNA fragmentation in *D. melanogaster* testis or ovaries, respectfully.

By incorporating natural population data of *D. yakuba* and *D. simulans* (Data S5) we polarized the SFS of each tRNA gene with 20kb flanking regions to test for selective sweeps at tRNA loci through statistical tests of neutrality. We generated null SFS distributions by the method of Hudson (*53*) with 10,000 simulations using a *D. melanogaster* population scaled mutation rate *θ* = 0.01328 (*54, 55*) and sought to identify departures from neutral expectations by applying Tajima’s D and Fay-Wu’s H tests to the polarized SFS. For all window and step configurations tested we were unable to reject the null hypothesis (*p* > 0.05) of the D and H tests, indicating non-directional variations in allele frequencies at tRNA loci. As the strongly conserved structure and short sequence of tRNAs may limit the power of spectra-based tests, we next conducted a Hudson-Kreitman-Aguadé-like (HKA-like) test to assess whether tRNA loci exhibited inter- and intra-specific site variability consistent with natural selection (*56*). We observed 36 tRNA genes with significant ratios of polymorphic to interspecific divergent sites compared to neutral expectations across all window and step configurations considered (Data S6). These represented 17 isotypes and gene ages ranging from B-1 to B7 (Fig. 4B, C). In total, four tRNA genes were both differentially expressed in gonads and had significant (*p* ≤ 0.05) HKA ratios. Strikingly, two of those four were *tRNA-Lys-TTT-2-5* and *tRNA-Asp-GTC-3-1* which we previously identified as the youngest differentially expressed tRNAs in *D. melanogaster* gonads (Fig. 4D). Further, when scrutinizing proportions of codon usage in *D. melanogaster* considering our differential expression and population genetic analyses, we found that *tRNA-Lys-TTT-2-5* and *tRNA-Asp-GTC-3-1* are the only two isoacceptors which correspond to the preferred codon of their isotype, have signatures of selection, and are differentially expressed between reproductive tissues (Fig 4E). Despite these findings, TTT is not the most abundant anticodon of tRNA^Lys^ isoacceptors by copy number. Of the remaining 36 tRNA genes with signatures of selection, six correspond to a major codon while four conserved isodecoders: B0 gene *tRNA-Ala-AGC-2-11* and B-1 genes *tRNA-Ala-AGC-2-10, tRNA-Ala-AGC-2-9, tRNA-Ala-AGC-2-8* are also differentially expressed. While we observed sexually dimorphic fragmentation patterns of tRNAs from 32 isoaccepting groups (Data S4), 13 were fragments of tRNA genes at genomic regions under directional selection, and only six in testis and 12 in ovary were derived from tRNAs corresponding to the preferred codon of their isotype. Most of the *D. melanogaster* tRNA repertoire (221/291) is neither differentially expressed between male and female reproductive tissues nor within genomic regions associated with non-neutral allele frequency variation. These data suggest tRNA repertoire composition can rapidly fluctuate while some copies are maintained in metazoan populations through selection of sex-associated function.

### Substitutions related to structure, function, and age suggest evolution of functionality

The paucity of high frequency minor alleles observed at tRNA loci may be attributable to the structure of tRNA which is highly conserved and specialized (Fig. 5A). We hypothesized that inferred gene age or functionally relevant structures of tRNA may affect SNP frequencies. To this end, we standardized base positions across *D. melanogaster* tRNA genes of varying length by aligning them to a covariance model of eukaryotic tRNA structure with INFERNAL v.1.1 (*57*). SNPs were observed to be present in all structural features of tRNA with the fewest in the anticodon loop and the majority in the acceptor stem (Fig. 5B), reflecting a functional constraint on SNP location. Loop regions contained fewer SNPs than stems highlighting their functional importance for aminoacylation as well as ribosome and aaRS recognition, suggesting purifying selection against SNPs within loops specifically (*58*). We next tested whether gene age correlated with SNP frequency and location (Fig. S4). We observed no significant relationships (*p* > 0.05, Pearson’s correlation) between inferred gene age and mean number of SNPs in each of the structural features of tRNAs (Fig. 5C, Data S7). These results indicate that tRNA copies originating at different time points experience equivalent purifying selection to reduce deleterious mutations.

**Fig. 5.**
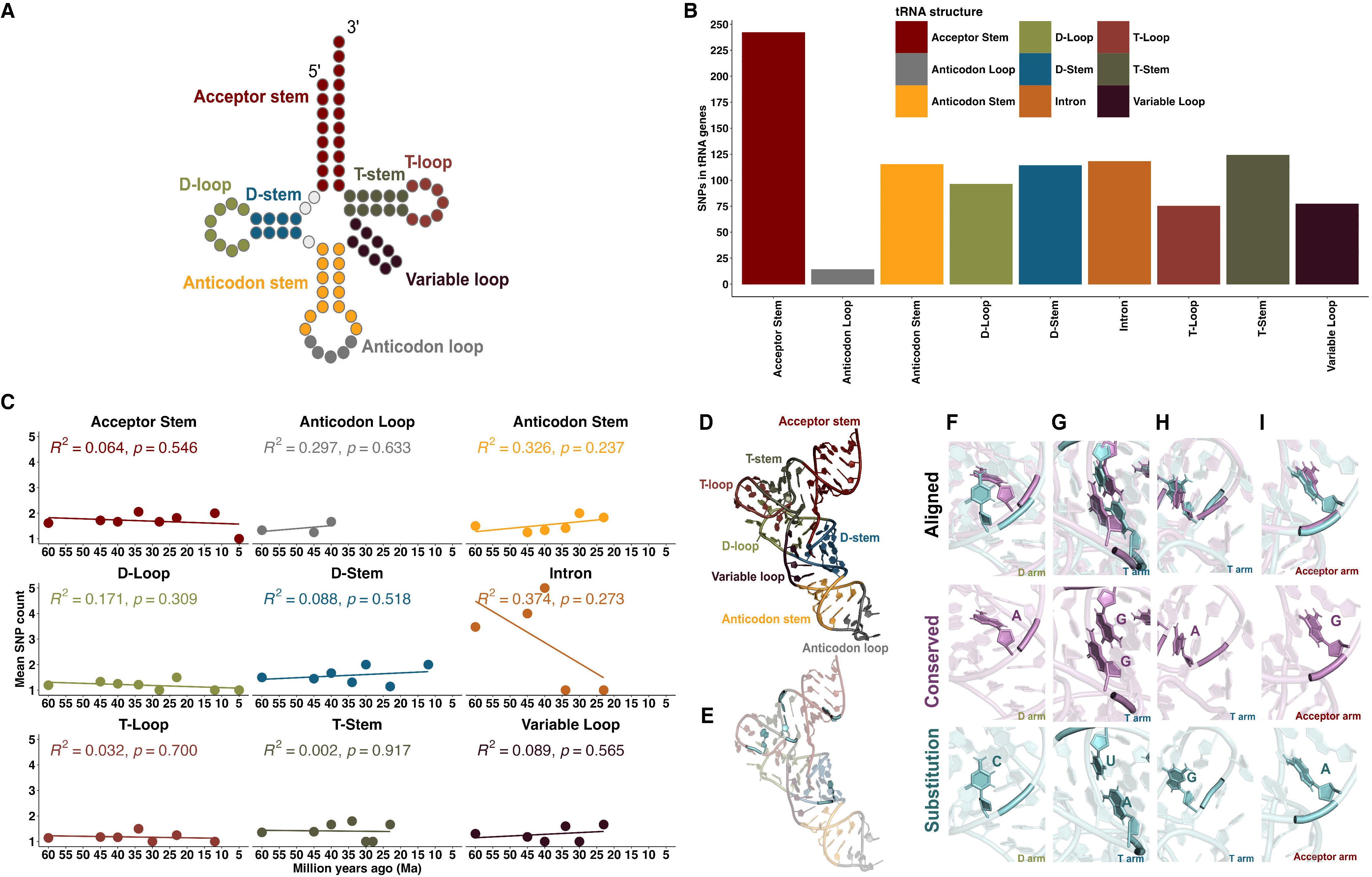
Mutation patterns are related to structure regardless of gene age. **(A)** Illustration of tRNA secondary structure. **(B)** Distribution of SNPs within structural features of tRNA. **(C)** Linear regression analyses illustrating the relationship between mean SNP density within structural features and the inferred evolutionary age of tRNAs given in million years (Ma). **(D)** Predicted tertiary structure of *tRNA-Glu-CTC-5-1* labeled with structural features. **(E)** Location of substitutions (blue) within *tRNA-Glu-CTC-5-1*. **(F-I)** Structural alignment (top row) of predicted tertiary structures (Data S7) of *tRNA-Glu-CTC-2-2* (purple, second row) and the *D. melanogaster*-specific isodecoder *tRNA-Glu-CTC-5-1* (blue, third row).

Following our observation of similar mutation patterns between tRNA duplicates, we next asked how fixed substitutions in taxon-specific tRNA genes may affect structural integrity and stability. Despite the recent inferred origination of the *melanogaster*-specific isodecoder identified previously, *tRNA-Glu-CTC-5-1,* it has five fixed substitutions compared to syntenic orthologs (Fig. 5D,E) and we estimated a substitution rate of 0.0597*bp*^−1^*Ma*^−1^in this gene using orthologous tRNAs from species spanning B3-B8 (*59*). We aligned the predicted structures (Data S8) of *tRNA-Glu-CTC-5-1* to the nearest conserved Glu isoacceptor, B4 gene *tRNA-Glu-CTC-2-2.* The transversion within the taxon-specific *tRNA-Glu-CTC-5-1* at residue C24 disrupted Watson-Crick base pairing of A24:U11, one of several identity determinants for glutamylation of tRNA^Glu^, potentially allowing a tertiary bond between residues 9 and 10 in the D-loop (*60*) (Fig. 5F). Two substitutions at positions 50 and 63 permitted Watson-Crick base pairing of 50U:63A, eliminating compensatory Hoogsteen interactions of 50G:62C and 63G:49C observed in *tRNA-Glu-CTC-2-2* (Fig. 5G). The substitution at 57G in the U-turn motif of the T-loop may stabilize tertiary interactions between the D- and T-loops (*61*), and the orientation of the transition at residue 68A at the 3’ end of the acceptor stem is flipped compared to the conserved 68G, potentially disrupting base pairing with 4C (Fig. 5H-I). These data illustrate that despite strong purifying selection and largely a lack of signatures of directional selection, recently duplicated tRNAs can rapidly accumulate substitutions yielding novel isodecoders (Fig. S5).

## DISCUSSION

Coordination of cytosolic tRNA availability and translational demand is essential to Darwinian fitness and cellular functionality across all domains of life (*62, 63*). Yet, our results show that the supply of translational machinery is neither conserved between species nor optimized in copy number to solely match global translational requirements. Rather, our work suggests the existence of translational conflict between unbiased codon usage of new protein coding genes and the nuclear encoded tRNA repertoires of metazoans which themselves expand and contract in copy number constantly over evolutionary time. Taxonomically restricted and isodecoding tRNA duplicates originating in recently diverged Drosophilid species exhibit patterns of sexually dimorphic functions and characteristics such as sex-biased abundance, fragmentation, and modification. The preservation of seemingly redundant tRNA gene copies is likely influenced by population genetics, nearly neutral processes, and sexual selection––not solely environmental factors (e.g., nutrient-driven tRNA modification) or directional selection for translational efficiency and accuracy, particularly at the genome-wide level (*64, 65*).

These findings expand our understanding of how origination time, nearly neutral processes, and sex differences affect the evolution and function of essential non-coding RNA. Prior to this work, mRNA decoding efficiency optimized through global adaptive codon usage was primarily invoked to explain the plurality of tRNA genes encoded by metazoan genomes (*66, 67*). The results reported here demonstrate that genetic drift or the acquisition of sexually dimorphic expression and fragmentation can influence the fate of new tRNA duplicates. Thus, in addition to multiple tRNA copies supporting translational efficiency, new isodecoding and isoaccepting tRNA copies and their fragments can contribute to maximizing the fitness of each sex.

## Acknowledgments

We thank members of the Long and Pan laboratories, N. Peña, A. K. Puppala, and M. Steinrücken for valuable discussion and comments on this work. We also thank M. Ludwig for generously sharing *Drosophila* stocks.

## Funding

National Institutes of Health T32 GM007197 Genetics & Regulation (DS)

National Institutes of Health T32 GM139782 Genetic Mechanisms & Evolution (DS)

National Institutes of Health 1R01GM116113-01A1 (ML)

## Author contributions

Conceptualization: DS, ML

Methodology: DS, MS, JC, SX, TP

Investigation: DS, MS, JC, SX

Visualization: DS

Funding acquisition: DS, ML, TP

Writing – original draft: DS

Writing – review & editing: DS, MS, JC, SX, TP, ML

## Data and materials availability

All data and software necessary to replicate analyses and figures presented in this manuscript are available at Zenodo (*68*), the main text, or supplementary materials

