## supplement for "Sex-biased duplicates are rapidly generated during *Drosophila* tRNA repertoire evolution"

**The PDF file includes:**

Materials and Methods
Figs. S1 to S4
Table S1

**Other Supplementary Materials for this manuscript include the following:**

Data S1 to S8

### Materials and Methods

#### Biased synonymous codon usage and tRNA adaptation test

Coding sequence annotations for D. melanogaster were retrieved from NCBI release 6.57. Frequencies of codon usage per gene were computed with codonw v.1.4.4 (*69*).

Gene-wide biased codon usage was compared using the translational adaptation index (tAI) as described in (*70*). Statistical analysis and visualization of results were performed with custom R scripts (*71*).

$$W_i = \sum_{j=1}^{n_i} (1 - s_{ij}) tGCN_{ij}$$

$$w_i = \begin{cases} \frac{W_i}{W_{max}} & \text{if } W_i \neq 0 \\ w_{mean} & \text{else} \end{cases}$$

$$tAI_g = \left(\prod_{i=1}^{l_g} w_{i_{kg}}\right)^{\frac{1}{l_g}}$$

$$f(x_g) = a + x_g + \frac{b}{{x_g}^2 (c - x_g)^2}$$

$$Nc_g = f(x_g) - \phi_g + \epsilon_g$$

$$\psi_g = f(x_g) - Nc_g$$

Briefly, $W_i$ represents the absolute adaptiveness value given as the sum of selective constraints on codon-anticodon coupling $s_{i,j}$ multiplied by tRNA gene copy number, $tGCN_{ij}$, for the $j_{th}$ tRNA that recognizes the $i_{th}$ codon to the $n_{th}$ isoacceptor that recognizes the $i_{th}$ codon, excluding stops. Relative adaptiveness values of codons are given by $w_i$ given by $W_i/W_{max}$ if the codon's absolute adaptiveness is not equal to 0, or the geometric mean of all $w_i$ with $W_i \neq 0$ otherwise. Adaptation of a gene to the genomic tRNA pool is represented by $tAI_g$ as the geometric mean of the relative adaptiveness values of $k$ codons along the length $l$ of a gene $g$. Effective number of codons $N_c$ of a gene $g$ were computed using an optimized version of Wright's equation, $f(x_g)$, (*72*) as described in (*70*). Estimates of translational selection for biased synonymous codon usage was quantified as the difference between the expected $N_c$ under neutrality and observed $N_{cg}$. The non-parametric correlation $S$ represents co-adaption between $\psi_g$ and $tAI_g$ which is a single value between -1 and 1 where high correlation coefficients indicate stronger translational selection.

#### Genomic sequence retrieval and annotation

24 Drosophilid long-read assemblies and their protein coding gene annotations were downloaded from NCBI. For each of the 24 assemblies we annotated tRNA genes using

tRNAscan-SE v.2.0.9 (*29*), discarded predicted pseudogenes, and kept only a set of high confidence annotations for downstream analyses.

**Gene age dating**

A Drosophilid 24-species divergence time estimated phylogeny was obtained from TimeTree (*31*). For each of the 291 D. melanogaster tRNA genes we first identified its two nearest protein coding genes flanking each side to create a set of four protein coding “anchor” genes. These anchor genes delineated a region containing one or more tRNA genes flanked by the anchors. By repeating this procedure, we identified tRNA and anchor gene sets in the 23 outgroup Drosophilid species which together with the D. melanogaster data were used to search for orthologous tRNA genes without convolution by extraneous, non-syntenic sequences. Our method largely reduced the homology search space and the probability of incorrectly aligning tRNA genes which would lead to over- or underestimation of gene ages by determining syntenic regions between conserved protein coding genes. For each D. melanogaster tRNA and anchor gene set, we identified orthologs and extracted their genomic region between their two anchor genes. We then conducted local alignments using the extracted outgroup anchor region and a given D. melanogaster tRNA gene to determine ortholog relationships (*73*). With the resulting alignments D. melanogaster tRNA gene ages were assigned according to a parsimony-based strategy similar to the methodologies of Zhang et. al. (*23*) and Zhou et. al. (*74*). Briefly, the age of a D. melanogaster tRNA gene was inferred based on the presence or absence of syntenic orthologs in the other 23 Drosophilid species. If a tRNA gene had orthologs in multiple species, the gene age was assigned to the phylogenetic branch of the oldest species with an alignment length of at least 70bp to avoid underestimation of gene ages. We further conducted a branch-wise analysis of tRNA gene presence and absence to identify tRNA loss events throughout the Drosophila phylogeny. If a gene was present in species on branch X but lacks a syntenic ortholog in any of the species on the next branch or younger, the gene would be inferred as lost in a given species.

**Whole-genome variant calling**

Natural populations of D. melanogaster with whole genome short read resequencing (WGS) data were downloaded from NCBI SRA (*52, 75*). In total, we obtained 727 high-quality D. melanogaster WGS samples after filtering data with a sequencing coverage threshold of at least 10. The WGS mapping was based on the standard pipeline of GATKv4.1 (*76*). Briefly, we first conducted quality control, adapter trimming, and quality filtering for raw reads with FastQC v2 (*77*) and fastp (*78*). The cleaned reads for each sample were mapped to the dm6 reference genome using BWA v0.7.17 (*79*) with default parameters. Following the sorting, indexing, and marking of PCR duplicates, the variants were called with the pipeline of the Best Practice protocol of GATK v4.1 using HaplotypeCaller (*76*). All variants were then filtered based on the following threshold: quality score of 30, sequencing coverage of 10x, genotype quality of 30, and a maximum of 25% missing data (*52*). The allele frequencies were analyzed with PLINK v1.9 (*80*). These data were used for downstream polymorphism analyses.

**Statistical tests of neutrality**

To test for natural selection at tRNA loci we first generated 10,000 neutral simulations following Hudson’s method (*53*) with 40,000 sites and 727 samples per simulation. For these simulations we used the following parameters: N0 = 1,000,000 (*81*), $\mu = 3.32 \times 10^{-9}$(*54*), and $\theta = 4 \cdot 1 \times 10^{6} \cdot 3.32 \times 10^{-9}$ where $\theta = 4N_0\mu$. We obtained empirical p-values for the Tajma’s

D and Fay-Wu's H statistics by identifying the number of simulated test values that were at least as extreme as observed values of D and H for all tRNA genes. We next polarized the D. melanogaster site frequency spectrum with population genetic data of D. simulans and D. yakuba to compute a Hudson-Kreitman-Aguadé-like test statistic. We used eight window and step configurations (400bp, 600bp, 800bp, 1kb and 25, 50bp) to collect observed polymorphic and divergent sites for comparison to neutral expectations (*82*). Only those tRNA loci that had significant HKA-like results in all window and step combinations were considered significant for downstream analysis.

**Structural analysis**

Tertiary tRNA structures were predicted with trRosettaRNA (*83*) and structurally aligned with US-align (*84*). Visualization was performed with PyMol (*85*).

**tRNA extraction, sequencing, and analyses by the MSR-seq pipeline**

Total RNA was extracted in biological triplicate from adult Iso-1 D. melanogaster adult reproductive tissues (ovary, testis) using TRIzol (ThermoFisher cat. no. 15596026) following manufacturer instructions. Sequencing libraries were prepared using the multiplexed small RNA (MSR) protocol of Katanski et. al. and sequenced on Illumina platform (*40*). Read alignment, quality control, abundance estimation, and statistical analyses were conducted using the tRAX software package (*86*).

**Tissue specificity analysis**

The tissue specificity metric spm (*49, 87*) was computed using the log-transformed expression matrix obtained from the small RNA sequencing analysis of all tRNA reads.

$$X = (x_1, x_2, \dots, x_i, \dots, x_{n-1}, x_n)$$

$$SPM_i = \frac{x_i^2}{|x_i| * |X|}$$

Where X is a vector of expression values of a tissue 1 through n, the number of samples. SPM values range from 0 to 1, where larger values indicate more specific expression, and values ≥0.9 are considered the most biased.

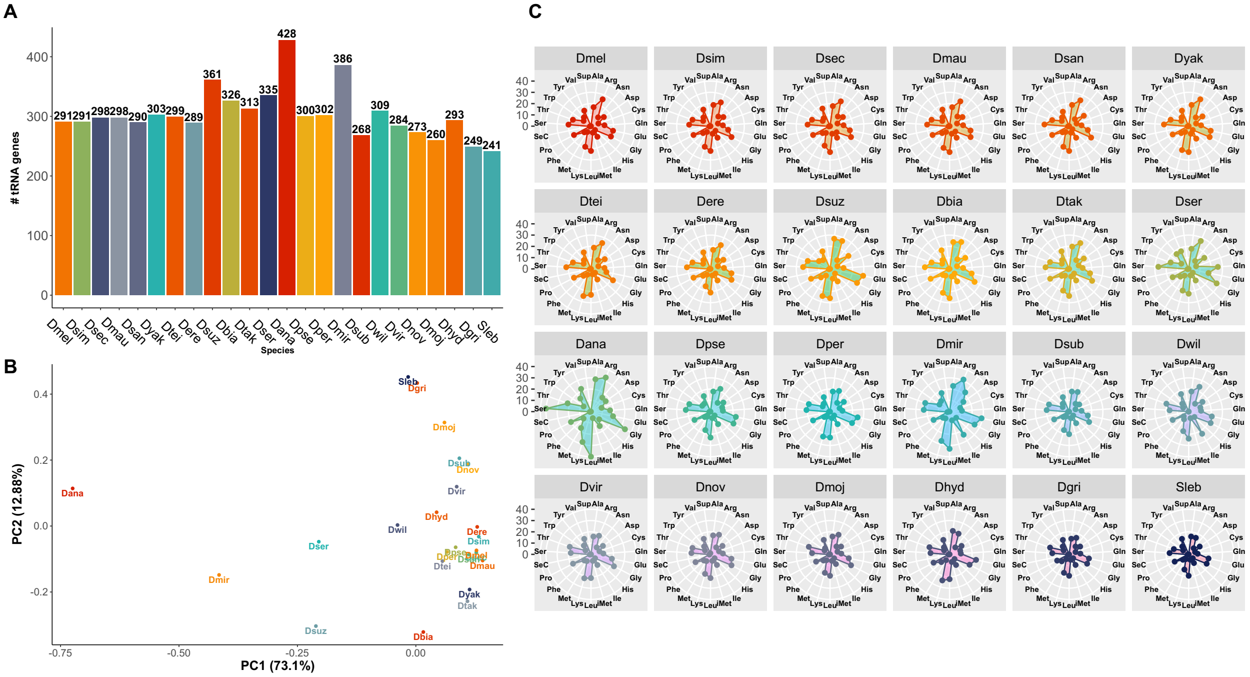


**Fig. S1. tRNA copy number diversity between Drosophilids.** (A) Total tRNA copy numbers per species considered in this work. Ordered from left to right by increasing time of divergence (rightmost, Sleb, is outgroup). (B) Principal component analysis of isotype and copy number variation in 24 Drosophilids. (C) Radar plots illustrating isotype copy numbers in each species, ordered left to right by row in order of increasing time of divergence. Each white ring indicates an increase of 10 copies, marked on the y-axis 0 through 40.

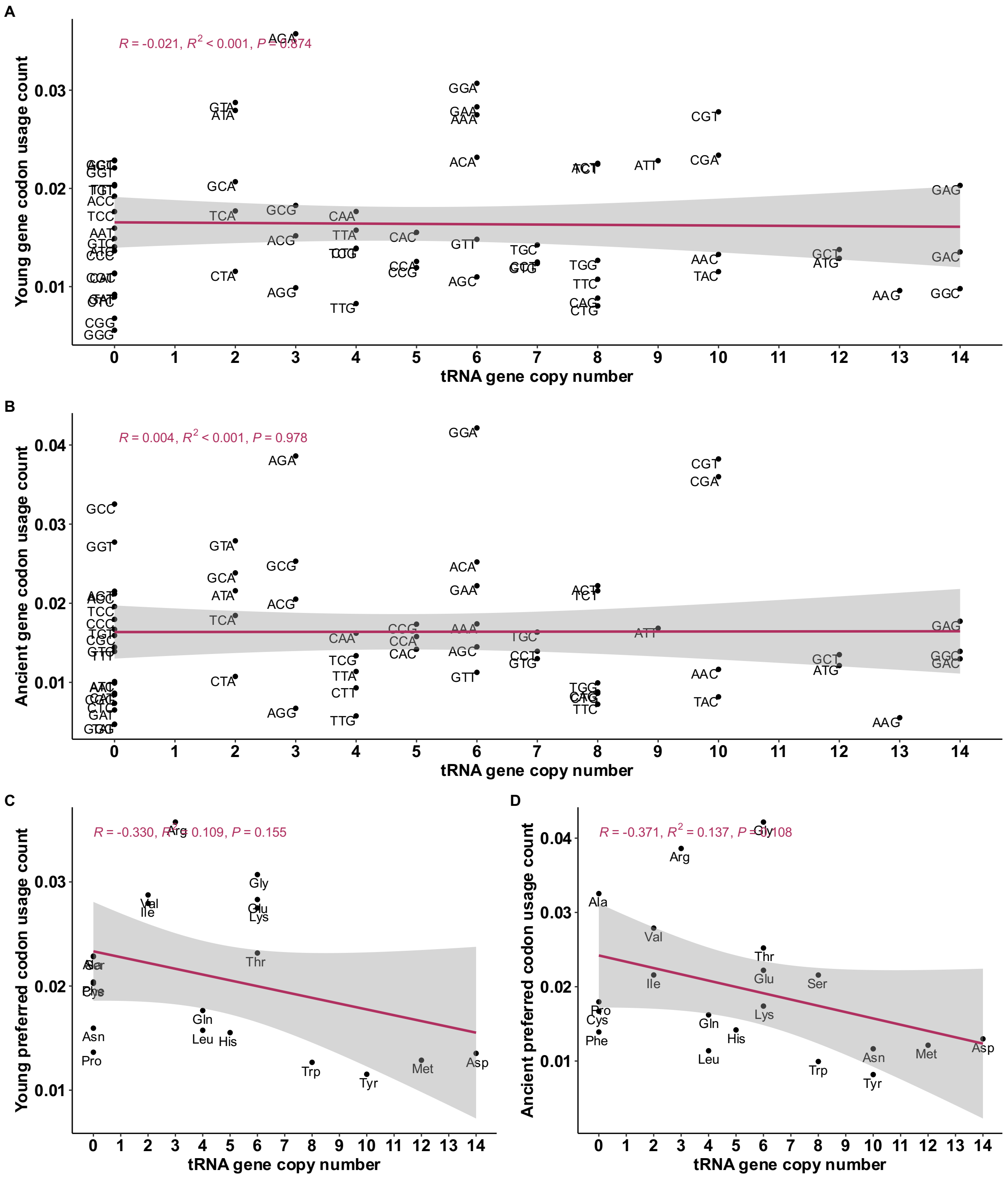


**Fig. S2. Increased tRNA copy number does not necessarily indicate increased usage of the corresponding codon.** (A) Young and (B) ancient genes show no significant correlation between tRNA isotype copy number and codon usage frequency (p=0.155 and p=0.987, Pearson's correlation). (C) Young and (D) ancient preferred codons per isotype are negatively correlated (p=0.013 and p=0.050, Pearson's correlation) with tRNA isotype copy number (*69*).

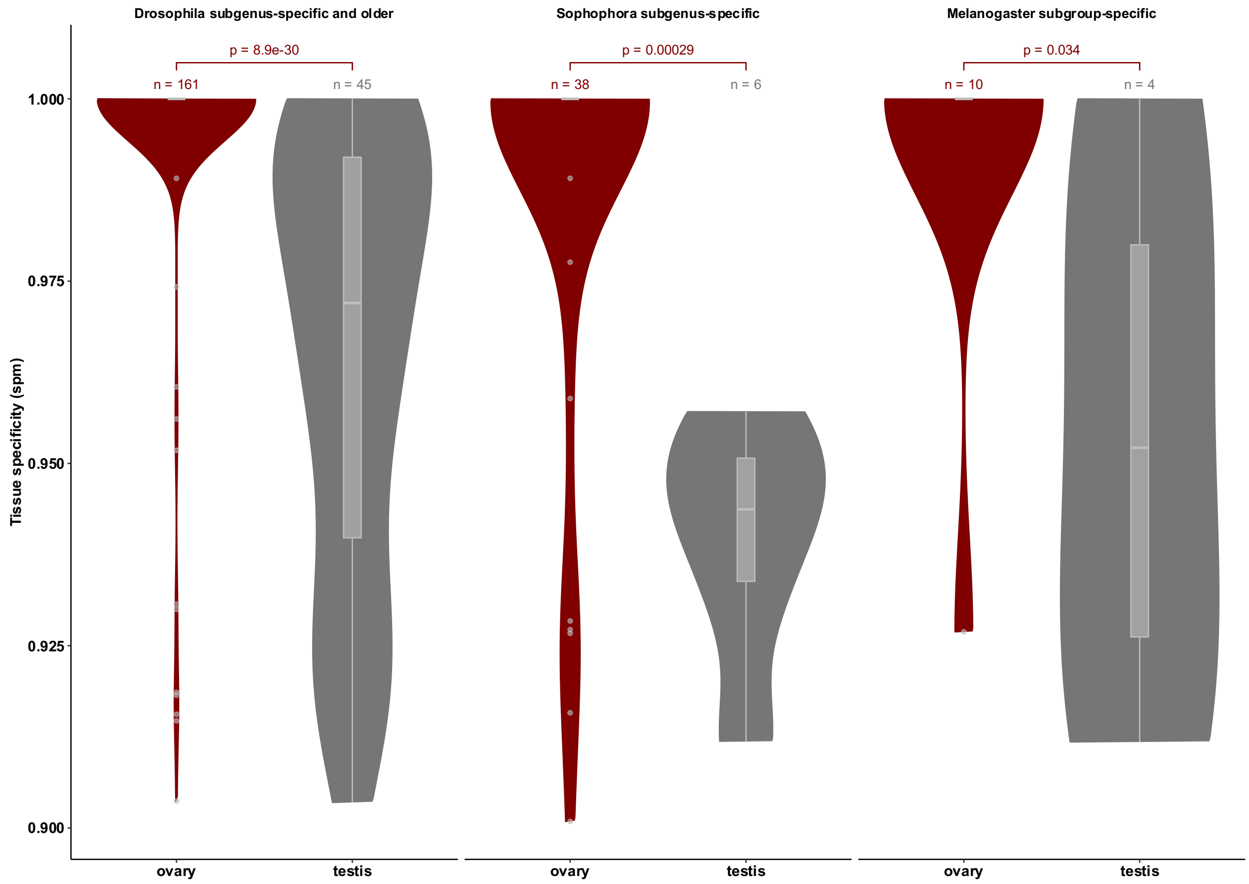


**Fig. S3. Ovarian tissue has more strongly tissue-biased tRNA transcripts than testis**. Distribution of the tissue specificity measure (spm) values ≥ 0.9 within ovary and testis of tRNA transcripts which we inferred to have duplicated within the Drosophila subgenus, Sophopohora subgenus, or melanogaster subgroup.

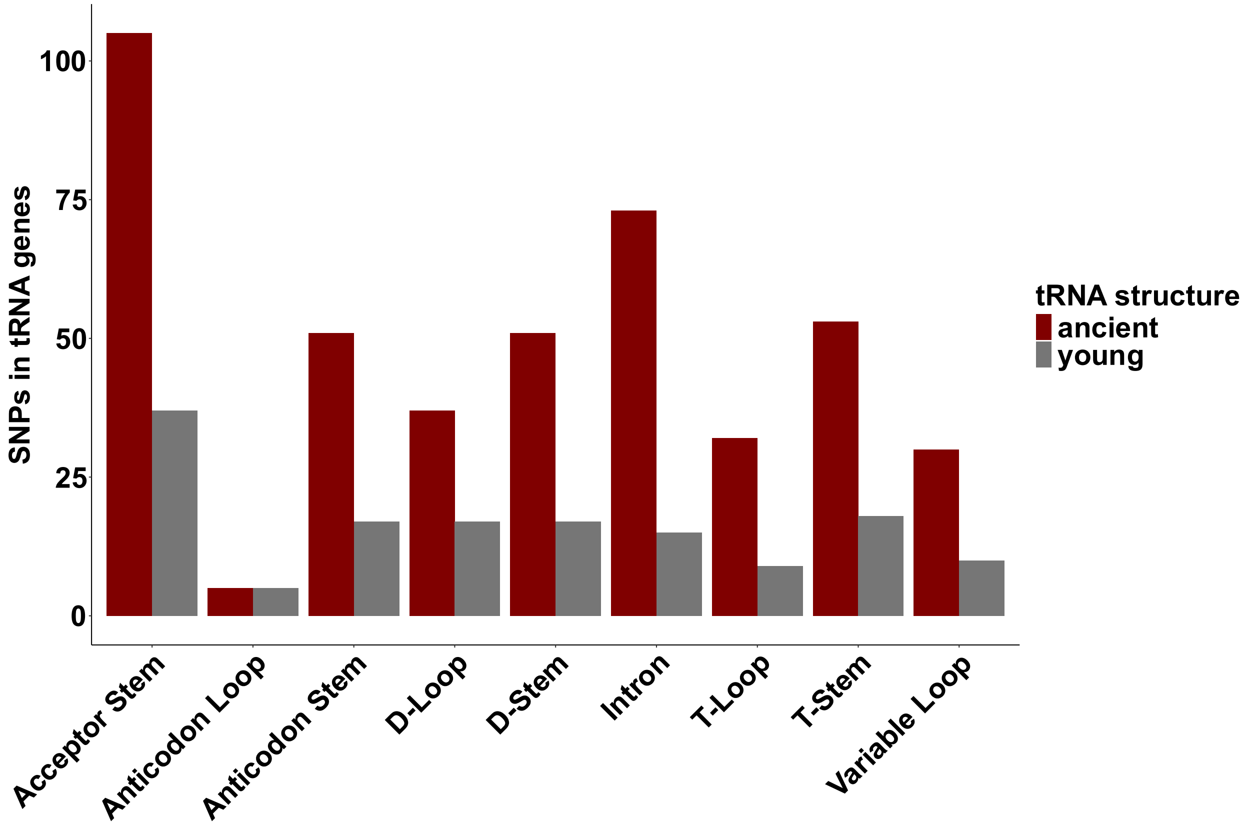


**Fig. S4. Distribution of SNPs within tRNA tertiary structure.** Total individual SNPs found within structures of 222 tRNA genes separated into ancient and young groups.

**Table S1.** List of genome assembly accessions used in the study.

| Species | Accession | Technology |
|---|---|---|
| Scaptodrosophila lebanonensis | GCF_003285725.1 | PacBio, long-read |
| Drosophila melanogaster | GCF_000001215.4 | PacBio, long-read |
| Drosophila serrata | GCF_002093755.2 | PacBio, long-read |
| Drosophila virilis | GCF_003285735.1 | PacBio, long-read |
| Drosophila novamexicana | GCF_003285875.2 | PacBio, long-read |
| Drosophila hydei | GCF_003285905.1 | PacBio, long-read |
| Drosophila persimilis | GCF_003286085.1 | PacBio, long-read |
| Drosophila erecta | GCF_003286155.1 | PacBio, long-read |
| Drosophila miranda | GCF_003369915.1 | PacBio, long-read |
| Drosophila mauritiana | GCF_004382145.1 | PacBio, long-read |
| Drosophla sechellia | GCF_004382195.2 | PacBio, long-read |
| Drosophila subobscura | GCF_008121235.1 | PacBio, long-read |
| Drosophila pseudoobscura | GCF_009870125.1 | PacBio, long-read |
| Drosophila suzukii | GCF_013340165.1 | PacBio, long-read |
| Drosophila teissieri | GCF_016746235.2 | PacBio, long-read |
| Drosophila santomea | GCF_016746245.2 | PacBio, long-read |
| Drosophila yakuba | GCF_016746365.2 | PacBio, long-read |
| Drosophila simulans | GCF_016746395.2 | PacBio, long-read |
| Drosophila ananassae | GCF_017639315.1 | PacBio, long-read |
| Drosophila takahashii | GCF_018152695.1 | PacBio, long-read |
| Drosophila grimshawi | GCF_018153295.1 | PacBio, long-read |
| Drosophila mojavensis | GCF_018153725.1 | PacBio, long-read |
| Drosophila willistoni | GCF_018902025.1 | PacBio, long-read |
| Drosophila biarmipes | GCF_025231255.1 | PacBio, long-read |

**Data S1 (separate file).**
D. melanogaster tRNA age dating annotations

**Data S2 (separate file).**
tRNA gene not lost during Drosophilid evolution.

**Data S3 (separate file).**
Differentially expressed tRNA genes.

**Data S4 (separate file).**
Sex-biased tRNA derived fragments.

**Data S5 (separate file).**
742 natural Drosophila population sample accessions.

**Data S6 (separate file).**
D. melanogaster tRNA genes with significant HKA-like test results.

**Data S7 (separate file).**
Location of SNPs within tRNA gene structural features.

**Data S8 (separate file).**
Structural alignment of tRNA-Glu-CTC-5-1 and tRNA-Glu-CTC-2-2.
